# Winter metabolic compensation endangers woodland salamanders under climate change

**DOI:** 10.1101/2025.10.11.681837

**Authors:** Braulio A. Assis, Isabella J. Burger, Savannah J. Weaver, Karey E. Duvall, Eric Riddell

## Abstract

Understanding the mechanisms that shape species’ geographic distributions is essential for predicting responses to climate change, particularly under winter conditions, which are warming more rapidly than summer conditions in many temperate ecosystems. Woodland salamanders, which constitute many of the species in the global hotspot of salamander diversity located in the southern Appalachian Mountains of North America, are thought to cope with harsh winters through dormancy and metabolic suppression. We used a field-based experiment to investigate the winter physiology and energetics of a woodland salamander (*Plethodon metcalfi*). Despite showing histological signs of dormancy, salamanders exhibited higher metabolic rates during winter than the active season when measured at low temperatures associated with winter, consistent with metabolic compensation. High-elevation salamanders also exhibited elevated metabolic rates at cold temperatures compared to low-elevation salamanders, indicative of countergradient metabolic compensation. When integrated into a biophysical species distribution model, depletion of energy stores estimated from winter metabolic rates under current climates accurately predicted the geographic range limits of our focal species. Under future warming scenarios, incorporating winter physiology indicated that 77% of the species’ range would experience local extinction due to depletion of energy stores. These findings challenge assumptions about winter energetics, reveal metabolic limits on cold-season survival, and highlight metabolic demands during dormancy as a key constraint on responses to climate change.

## Introduction

Organisms experience variable environments across spatial and temporal scales (Lawson et al., 2015), and in response, they adjust physiological traits to facilitate persistence across geographic gradients and seasonal cycles (Norin & Metcalfe, 2019). As a key determinant of energy balance, survival, and reproduction, plasticity in metabolic rates plays a central role in facilitating responses to environmental variation across space and time (Burton et al., 2011; van der Meer, 2006). For instance, metabolic suppression allows for energy conservation in response to high temperatures or long periods of inactivity (Storey & Storey, 2004). Conversely, metabolic compensation – an increase in metabolic rates at cool temperatures – allows organisms to maintain physiological function despite the inhibitory effects of low temperatures on biochemical reaction rates (Catenazzi, 2016; Williams et al., 2014). However, the capacity of organisms to adjust metabolic rates in response to rapid, novel environmental change, such as climate change, remains uncertain (Deutsch et al., 2015; Sun et al., 2022). Notably, most warming from climate change is expected to occur during the winter, with forecasts including reductions in winter duration, increased temperatures, increased number of freeze-thaw cycles, and even the loss of winter conditions altogether (Williams et al., 2015). Warmer winters are associated with reductions in body condition and survival, potentially driven by higher metabolic rates and greater energy expenditure (Reading, 2007). Investigating how metabolic rates vary across geographic and seasonal gradients can reveal how environmental factors shape physiology and constrain species’ persistence in changing climates over time.

Seasonal cycles drive predictable variation in temperature, precipitation, and resource availability in temperate ecosystems (Grant et al., 2017; Yanco et al., 2022). Organisms have evolved suites of strategies to optimize fitness in response to these recurring environmental changes (McKechnie et al., 2015; Scherbarth & Steinlechner, 2010). To cope with unfavorable conditions, such as cold winter temperatures and scarce resources, many temperate species enter periods of metabolic inactivity, such as hibernation or dormancy (Wilsterman et al., 2021). For ectotherms, cold temperatures are typically associated with low metabolic rates and constraints on activity, often triggering dormancy through further metabolic suppression to conserve energy (Lailvaux, 2007; Storey & Storey, 2004). During periods of dormancy, animals undergo significant morphological changes in tissues and organs, including reductions in intestinal mass, intestinal length, mucosa thickness, and villus length (do Nascimento et al., 2016; Hume et al., 2002), reflecting inactivity in nutrient acquisition and absorption (Goodrich et al., 2024). In amphibians, dormancy is also associated with increased epidermal thickness, possibly reflecting slowed epidermal turnover (Barni et al., 1987; Czopek, 1959; Kobelt & Linsenmair, 1986). The extent of these changes likely varies along environmental gradients, with factors like temperature influencing their expression and magnitude.

Metabolic responses often vary predictably across environmental gradients, with populations adapting through metabolic suppression or compensation (Conover et al., 2009). In ectotherms, individuals that live in warm climates (such as low latitudes or elevations) tend to suppress metabolic rate compared to populations from cool climates, minimizing the energetic costs imposed by higher ambient temperatures (Seebacher et al., 2015). Conversely, individuals that live in cool climates often exhibit elevated metabolic rates relative to individuals from warm climates, helping to offset reduced performance and slower growth rates at low temperatures. This specific pattern is described by the Metabolic Cold Adaptation hypothesis (MCA) (Clarke, 1993), which posits organisms adapt to cooler environments by increasing metabolic rate. More broadly, MCA represents a form of countergradient variation (Berven et al., 1979), where genetic influences on a trait oppose environmental effects, potentially leading to minimal geographic variation in the phenotype along environmental gradients (Conover & Schultz, 1995). Metabolic suppression or compensation are also critical strategies for responding to seasonal changes, helping organisms reduce the energetic costs of warm summers and boost physiological performance in the cool winters (Bullock, 1955; Catenazzi, 2016; Speers-Roesch et al., 2018). Unlike MCA, these seasonal responses involve reversible phenotypic plasticity within an individual, although the adaptive explanations are similar. For energetically limited species, these reversible plastic responses could be essential for persisting in the face of seasonal climatic extremes.

Woodland salamanders from the genus *Plethodon* provide an ideal system to study metabolic responses to environmental variation, as they are energetically constrained, experience seasonal fluctuations, and live along elevational gradients (Gifford & Kozak, 2012). Woodland salamanders are highly diverse and abundant in the temperate forests of the southern Appalachian Mountains in North America (Fig. 1A,B), a region recognized as the global hotspot of salamander diversity (Kozak et al., 2005; Kozak & Wiens, 2010a). These salamanders have also evolved an unusual and extreme strategy for conserving energy: the complete loss of lungs. This evolutionary loss is thought to reduce the energetic costs associated with respiration (Reagan and Verrell 1991) and contributes to woodland salamanders having one of the lowest metabolic rates among vertebrates (Feder et al., 1984). Still, their physiology during the winter months—when organisms are energetically limited—remains largely unstudied (Fraser, 1976; Gifford, 2016). Like other temperate amphibians, woodland salamanders are assumed to be inactive in the winter as individuals that emerge in the spring lack lipid reserves and can appear emaciated (Camp & Jensen, 2007; Fraser, 1976). However, studies have yet to establish whether woodland salamanders enter a period of physiological dormancy, and energy balance models have thus far ignored the potential effects of winter dormancy on habitat suitability under climate warming scenarios (Gifford and Kozak 2012, Riddell et al. 2018). Therefore, experimental studies on metabolic physiology may reveal fundamental mechanisms underlying overwintering responses and clarify how seasonal differences in metabolic physiology contribute to species’ vulnerability to climate change.

**Figure 1:**
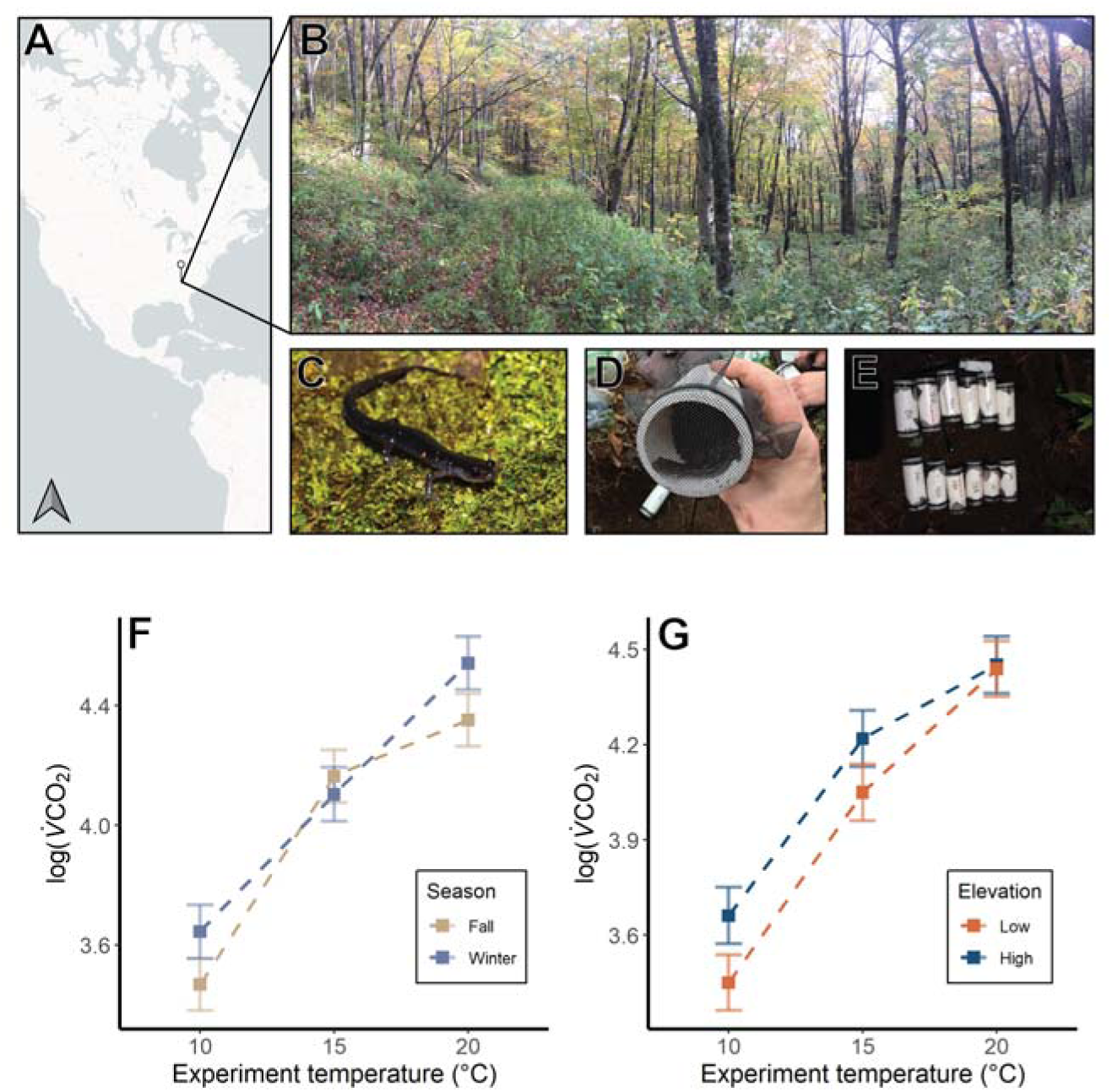
Evidence for winter metabolic compensation and countergradient variation in a woodland salamander (*P. metcalfi*). (A) A map of North America with a pin indicating the location of our study site. (B) The temperate forest in which the study was conducted. (C) The focal species, the southern gray-cheeked salamander (*P. metcalfi*). (D) A salamander inside the enclosure prior to being placed underground for the winter. (E) One hole containing the experimental enclosures prior to being buried with the HOBO datalogger. (F) Variation in V CO_2_ (μL h^-1^) as evidence of winter metabolic compensation at cold temperatures and fall suppression at warm temperatures illustrated by the interaction between experimental temperature and season. (G) Variation in V CO_2_ (μL h^-1^) as evidence for countergradient variation illustrated by the interaction between experimental temperature and elevation. High elevation salamanders had higher metabolic rates than low elevation salamanders, with differences more pronounced in colder temperatures. In both plots, points represent marginal means with 95% confidence intervals for a season-by-temperature interaction and for an elevation-by-temperature interaction controlling for the effects of sex, body mass, and either elevation (F) or season (G), along with individual identity as a random effect. Photos by Christopher Mayerl (C), EAR (B, D), SJW (E).

In this study, we conducted a field-based overwintering experiment on the metabolic physiology of a woodland salamander (*P. metcalfi*, Fig. 1C, D, E) to understand the energetic effects of winter warming on salamander physiology and distributions. Salamanders were placed in semi-natural field enclosures during the winter, then excavated mid-winter and early spring for laboratory respirometry and compared to salamanders collected directly from the field in the fall, using the volume carbon dioxide production (V CO_2_, μL h^-1^) as an index of metabolic rate. At the end of metabolic measurements, we also assessed histological markers of winter dormancy, termed brumation (Wilsterman et al. 2021), of the skin and digestive tissues. Within seasons, we also assessed metabolic variation in salamanders collected from high and low elevations to further evaluate whether salamanders suppress or elevate metabolism in response to naturally occurring temperature gradients across their elevational range. Our experimental design, which measured metabolic rates across a range of ecologically relevant temperatures (10, 15, 20 ), allowed us to distinguish between metabolic compensation and suppression, which are challenging to disentangle from measurements at a single temperature. We expected reduced metabolic rates at warmer temperatures to be indicative of suppression, whereas compensation would be indicated by increased metabolic rates at cooler temperatures. We predicted that salamanders, like many other ectotherms (Abe & Garcia, 1991; Boutilier et al., 1997; Podhajský & Gvoždík, 2016), would exhibit metabolic suppression during winter dormancy in an effort to reduce energy expenditure. Additionally, we hypothesized that low-elevation salamanders would exhibit lower metabolic rates than their high-elevation counterparts due to adaptation for metabolic suppression in their warmer environment. We then integrated the thermal sensitivity of metabolic rate into biophysically informed species distribution models to evaluate whether energy balance helps to explain current species’ range limits, determine whether depleting energy reserves leads to increased local extinction across the species’ range during the winter under climate change scenarios, and test for countergradient variation in metabolic rate across elevations. Together, these approaches provide a comprehensive understanding of how seasonal and elevational variation in metabolic physiology could mediate overwintering success and influence vulnerability to future climate change.

## Results

### Seasonal and elevational effects on metabolic rate

We observed significant variation in metabolic rates of salamanders across seasons, elevations of origin, and experimental temperatures. Metabolic rates were significantly influenced by the interactions of experimental temperature with season or with elevation (Table 1). Contrary to our expectations, winter salamanders exhibited higher metabolic rates than fall salamanders at 10°C (19.5%; post-hoc t-test, *t* = -2.78, *p* = 0.006) and 20°C (20.9%; *t* = -2.99, *p* = 0.003), but not at 15°C (Fig. 1F). The temperature × elevation interaction showed that high elevation salamanders had elevated metabolic rates at 10°C (23.7%, *t* = 3.32, *p* = 0.001) and 15°C (18.5%; *t* = 2.65, *p* = 0.009) compared to low elevation salamanders but not at 20°C (Fig. 1G). In addition, metabolic rates increased with body mass and were higher in males (Table 1).

**Table 1:** Type II ANCOVA table for predictors of log-transformed V CO_2_ to determine seasonal and elevational effects. *Elevation*: factor of 2 for high and low-elevation populations; *Mass*: body mass prior to respirometry assay; *Temperature*: factor of 3 for respirometry assays at 10, 15, and 20°C; *Season*: factor of 2 for individuals collected in the fall or winter; *Sex*: factor of 2 for males and females.

|  | Sum Sq | Mean Sq | DF | F | P |
| --- | --- | --- | --- | --- | --- |
| Elevation | 0.71 | 0.71 | 1 | 9.85 | 0.002 |
| Mass | 0.94 | 0.94 | 1 | 13.13 | <0.001 |
| Temperature | 33.60 | 16.80 | 2 | 223.31 | <0.001 |
| Season | 0.44 | 0.44 | 1 | 6.09 | 0.016 |
| Sex | 0.31 | 0.31 | 1 | 4.27 | 0.042 |
| Temperature:Season | 0.82 | 0.41 | 2 | 5.70 | 0.004 |
| Temperature:Elevation | 0.45 | 0.23 | 2 | 3.17 | 0.045 |

Due to sampling limitations for low-elevation populations in spring, we conducted a follow-up analysis focusing only on the high-elevation population, which was well-sampled across all three seasons (fall, winter, and spring). This analysis revealed a similar seasonal pattern (Table 2), with winter salamanders again exhibiting elevated metabolic rates at 10°C (*t* = -2.057, *p* = 0.041) and 20°C (*t* = -2.021, *p* = 0.045). Spring metabolic rates tended to be intermediate to fall and winter, suggesting a return to a state of activity or arousal (Fig. S3). Experimental temperature remained a significant predictor, but mass and sex were not (Table 2; Fig. S3). Of note, we did not observe a significant elevation × season interaction in any analysis, suggesting that metabolic compensation does not covary with elevation.

**Table 2:** Type II ANCOVA table for predictors of log-transformed V CO_2_ for seasonal analysis that included spring. *Mass*: body mass prior to respirometry assay; *Temperature*: factor of 3 for respirometry assays at 10, 15, and 20°C; *Season*: factor of 3 for individuals collected in the fall, winter, or spring; *Sex*: factor of 2 for males and females.

|  | Sum Sq | Mean Sq | DF | F | P |
| --- | --- | --- | --- | --- | --- |
| Mass | 0.02 | 0.02 | 1 | 0.26 | 0.61 |
| Season | 0.19 | 0.09 | 2 | 1.25 | 0.293 |
| Temperature | 20.53 | 10.27 | 2 | 134.98 | < 0.001 |
| Sex | 0.00 | 0.00 | 1 | 0.01 | 0.94 |
| Season:Temperature | 1.06 | 0.26 | 4 | 3.47 | 0.010 |

### Histological signatures of winter dormancy

To assess the physiological state of salamanders in winter, we performed histological analyses to identify morphological signatures of brumation. Salamanders showed significant differences in gut and skin morphology between seasons, consistent with the onset of brumation (Fig. 2, Table 3). Compared to fall salamanders, those sampled in the winter had 48.0% fewer intestinal villi (Fig. 2, *t* = 3.46, *p* = 0.004), their intestinal cross-sectional diameter was 23.7% narrower (Fig. 2, *t* = 2.67, *p* = 0.016), and their epidermis was 31.3% thicker (Fig. 2, *t* = -3.28, *p* = 0.001). Histological measurements from spring salamanders were either similar to those observed in winter or intermediate between winter and fall values (Fig. 2), suggesting that individuals were in a transitional physiological state—potentially shifting from dormancy to arousal. We observed no difference in the length of the intestinal villi between seasons (Table 3). Together, these histological markers indicate that salamanders were undergoing brumation in winter.

**Figure 2:**
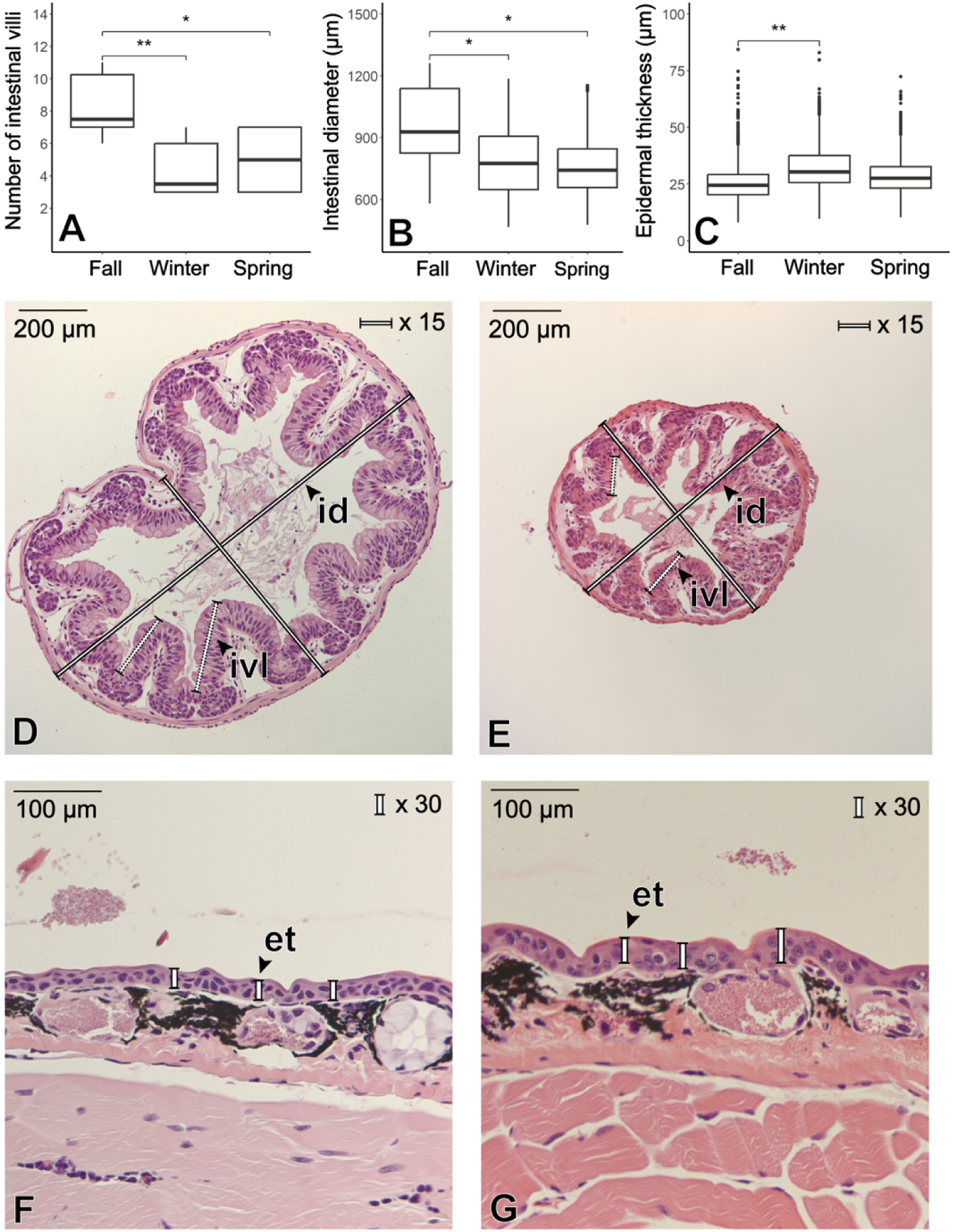
Histological evidence of winter dormancy in a woodland salamander. In relation to active salamanders in the fall, salamanders in winter had fewer intestinal villi (A), smaller intestinal diameter (B), and a thicker epidermis (C), consistent with dormancy. Boxplots represent the 25^th^, 50^th^, and 75^th^ percentiles, and whiskers represent the largest and lowest values within 1.5 times the interquartile range. Asterisks denote significant at α = 0.05 (*) and α = 0.01 (**). Histology images are cross sections of the lower intestine of a fall (D) and a winter salamander (E), and cross sections of the dorsal skin of a fall (F) and a winter (G) salamander. Bolded text indicates intestinal diameter (id), intestinal villus length (ivl), and epidermal thickness (et).

**Table 3:**
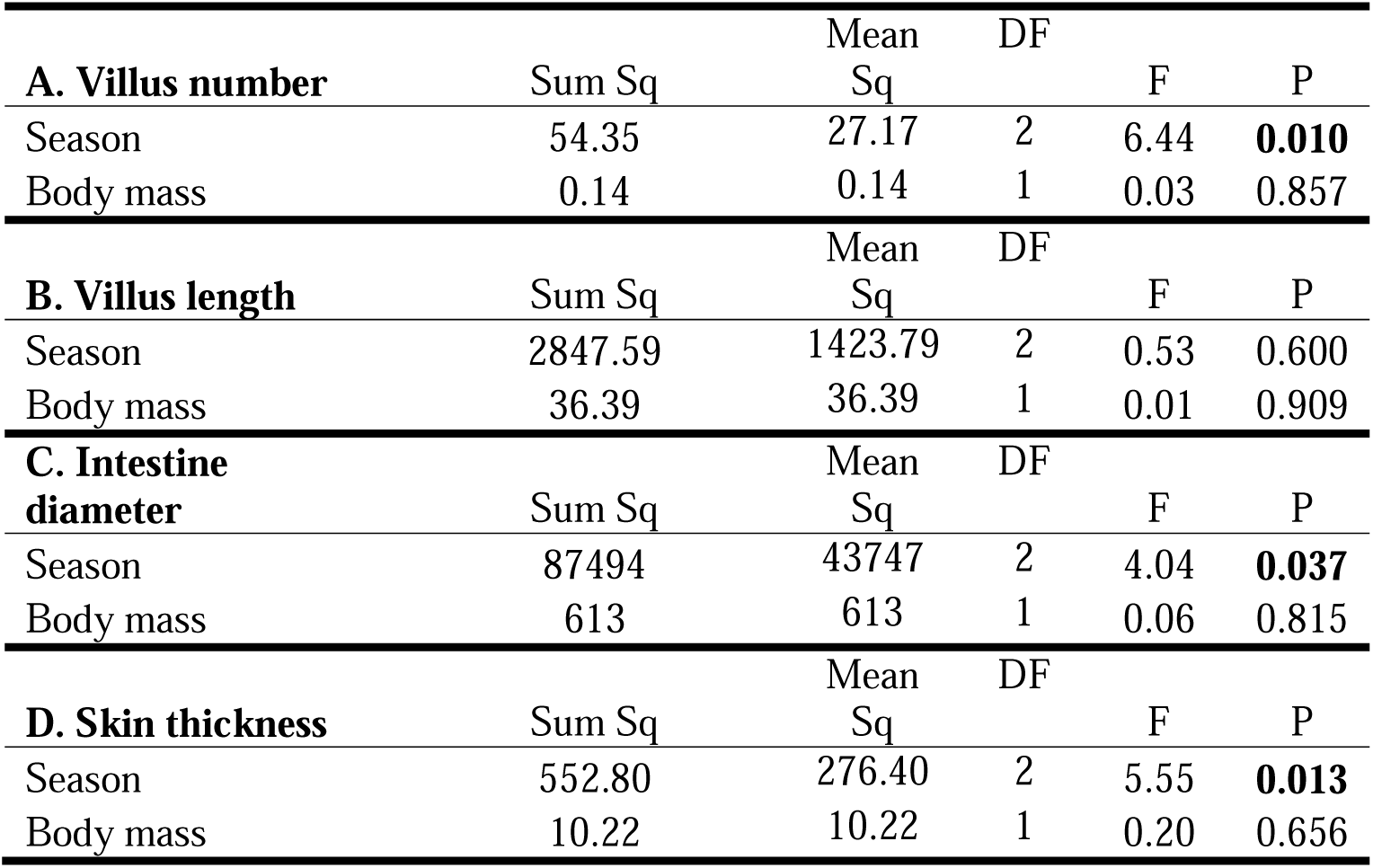
Type II ANCOVA table for a linear model (A) or linear mixed effects models for repeated measures within a single salamander (B, C, D) on morphological variation in lower intestine and skin of salamanders throughout active and inactive seasons. Season statistics are in relation to data obtained from salamanders in fall.

### Energetic costs in current and future climates

We used a biophysical species distribution model to investigate the impact of metabolic compensation during brumation on the range limits and extinction risk under a moderate future warming scenario. Our simulations suggest that winter energetics play a critical role in shaping the current range limits of *P. metcalfi*, as indicated by a negative correlation between the probability of presence and the proportion of total lipid reserves consumed across the species’ range (Fig. 3A,B). In the model, the probability of presence fell to zero at the point where salamanders were predicted to deplete their lipid reserves under current climate conditions (Fig. 3B). We also found that failing to account for metabolic compensation during winter dramatically underestimates energy use (Fig. 3D–F), and the observed metabolic compensation during winter under future warming scenarios might cause a substantial increase in local extinction across the range (Fig. 3F). Under current climate conditions, the model predicts that salamanders can sustain themselves through the winter using lipid reserves at nearly every site across their geographic range, regardless of whether fall or winter metabolic rates are assumed (proportion of sites in which salamanders deplete total lipid reserves: fall = 0.1%, winter = 11.8%). However, under future warming, the model predicts that salamanders at 26.0% of sites will experience local extinction by fully depleting their lipid reserves assuming fall metabolic rates, and at 77.0% of sites assuming the more relevant winter metabolic rates. Moreover, salamanders are predicted to consume substantially more —and in many cases nearly all—of their energy reserves during winter under future warming scenarios. Specifically, the model estimates that salamanders will use an average of 108.6 ± 11.9% of their available lipid stores during the winter across the geographic range under future warming scenarios, compared to 85.6 ± 9.2% under contemporary climate conditions. These percentages represent relative energy expenditure based on estimated lipid availability at the start of winter. Values exceeding 100% indicate that, under future warming, salamanders at many sites are predicted to consume more lipid reserves than they have available, suggesting a severe energetic deficit that could compromise overwinter survival.

**Figure 3.**
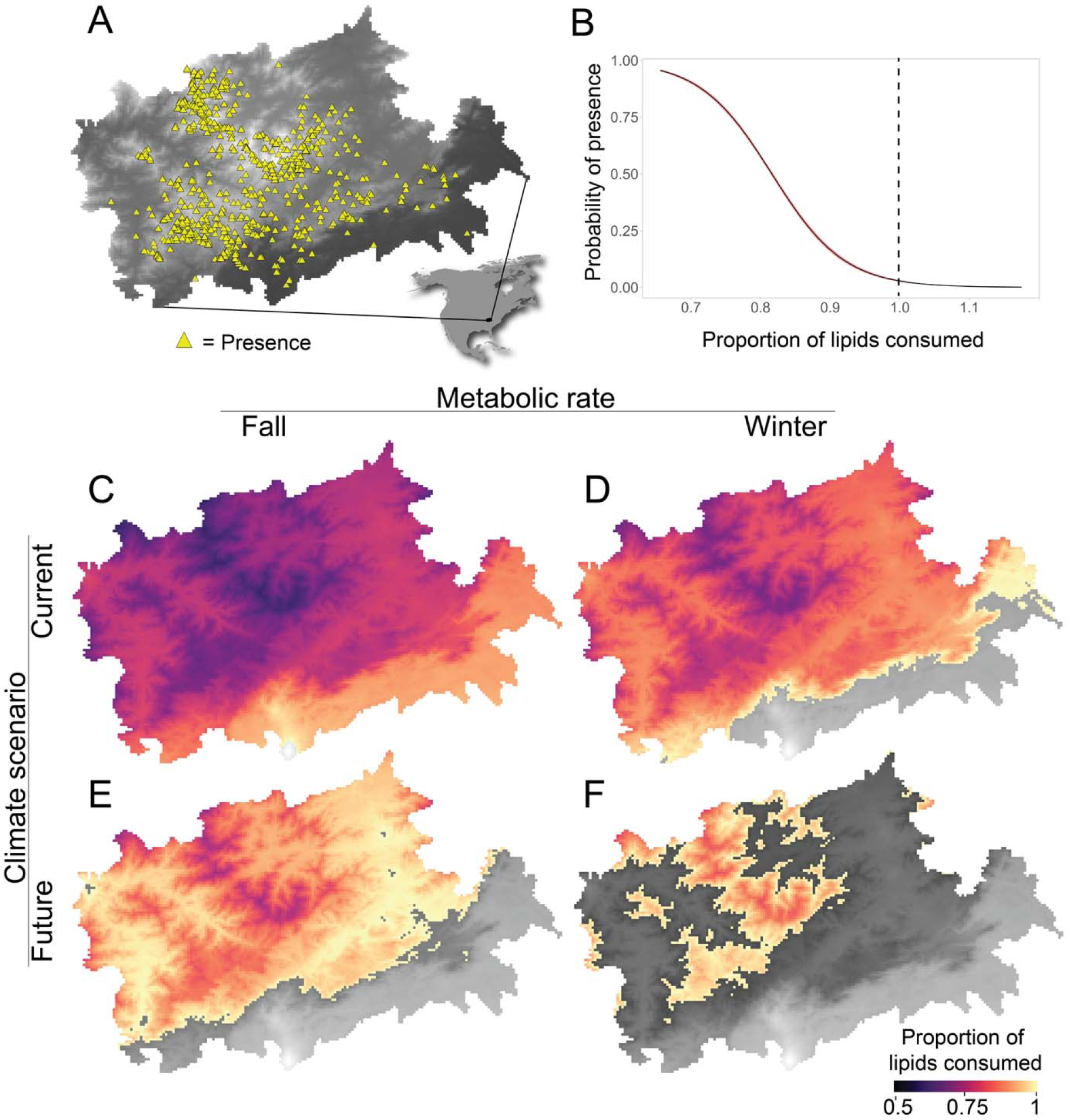
Greater erosion of suitable habitat under climate change due to winter metabolic compensation. (A) The geographic range of *P. metcalfi* in the Southeastern Appalachian Mountains with presence locations (yellow triangles) overlain on top of a digital elevation map. (B) The negative correlation between the probability of presence of *P. metcalfi* across their range with respect to proportion of lipids consumed during the winter. The probability of presence declines to nearly zero as the model predicts salamanders consume 100% of their lipid reserves. (C-F) Maps of proportion lipids consumed across the range of *P. metcalfi* assuming fall metabolic rates in contemporary climates (C), winter metabolic rates in contemporary climates (D), fall metabolic rates in future climates (E), and winter metabolic rates in future climates (F). In C-F, areas in gray correspond to 100% of lipid reserves consumed. Thus, models parameterized with winter metabolic rates exhibit the greatest reduction in regions that can support the energetic costs of winter, and salamanders are predicted to consume nearly 100% of their lipid reserves across the landscape under future climate change.

### *In silico* reciprocal transplant experiment

To test for countergradient variation in metabolic rate, we conducted an *in silico* reciprocal transplant experiment to estimate metabolic rates during the winter across elevations and thermal sensitivities. Specifically, we estimated metabolic rates underground in both a high-elevation and a low-elevation environment using the measured thermal sensitivity of metabolic rates of salamanders sourced from both elevations. This simulation revealed that phenotypic differences in winter energy expenditure were minimized when salamanders were modeled in their native elevation environments (Fig. S5). Specifically, low-elevation salamanders at low elevations and high-elevation salamanders at high elevations had the most similar metabolic rates. This pattern is consistent with countergradient variation, where genetic differences between populations offset environmental effects, leading to similar physiological performance across contrasting habitats. Although all pairwise comparisons were statistically significant (*p* < 0.001), these results were largely influenced by the large sample size (N = 25,044). Examination of effect sizes showed that most group comparisons had effect sizes that did not overlap zero, indicating meaningful differences. However, the comparison between low elevation salamanders at low elevations and high elevation salamanders at high elevations yielded an effect size overlapping zero (supplementary figure S5), with the lack of meaningful differences between these groups supporting countergradient variation.

## Discussion

We examined the metabolic physiology of a woodland salamander and found that, contrary to expectations, winter dormancy was marked by metabolic compensation at low temperatures, which they experience during the winter months. This response contrasts with our predictions and patterns observed across taxa, in which winter dormancy typically coincides with reduced energy expenditure (Dubiner et al., 2023; Podhajský & Gvoždík, 2016; Williams, Chick, et al., 2015). These findings are particularly surprising given the histological evidence that salamanders were in a state of dormancy, based on reduced intestinal diameter and number of intestinal villi and increased epidermal thickness. This analysis also indicated that salamanders in the fall suppress metabolic rates at warmer temperatures, revealing both metabolic compensation at cool temperatures in the winter and suppression at warm temperatures in the fall. Across elevations, salamanders from high elevations had higher metabolic rates than salamanders from low elevations when measured at cooler temperatures. The absence of metabolic differences at warmer temperatures suggests that these patterns are not driven by pressure for energy savings in warmer environments (Moffett et al., 2018). Rather, the elevational and seasonal patterns suggest that salamanders regularly use metabolic compensation to cope with the reduced capacity for homeostatic maintenance at low temperatures (Bullock, 1955; White et al., 2011). This work highlights a unique strategy among animals to elevate metabolic rates during winter dormancy— a period typically characterized by suppression—highlighting an important lower bound on the energetic requirements for maintaining homeostasis in low temperature environments. If such adaptive mechanisms are common but unaccounted for in bioenergetic models, they could result in unforeseen impacts on species distributions under climate change.

Our mechanistic species distribution model suggests that winter energetics impose strong constraints on species ranges, as shown in other species (Réveillon et al., 2019; Roberts & Williams, 2022). In our focal species, predicted depletion of energy reserves during winter dormancy closely matches observed patterns of presence across the landscape (Fig. 3). These results indicate that warmer winter temperatures, especially at low elevations, will increase metabolic costs during dormancy and drive local extinction due to the rapid depletion of energy reserves. Notably, some small, isolated southern populations currently persist in areas where the model predicted near-total depletion of lipid stores (Fig. 3D). This mismatch could reflect unmodeled microclimatic buffering (e.g., cool, humid riparian zones), local adaptation to reduce winter energy use, or plasticity in the degree of metabolic compensation (Hall et al., 2016; Pincebourde et al., 2016; Suggitt et al., 2015). Interestingly, the spatial constraints imposed by winter energetics parallel those previously identified for the active season, where limits on surface activity due to water loss were associated with the distribution of the same focal species (Riddell et al., 2017). These findings suggest that both energy limitations during dormancy and hydric constraints during the active months play critical roles in determining range boundaries. While these mechanisms could act synergistically, it is also possible that one is more dominant in shaping the species’ distribution. Disentangling their relative contributions will require targeted validation of how environmental conditions influence activity patterns and energy reserve depletion. Importantly, our results indicate that winter energy constraints might drive more extensive range contractions under climate change than expected for active-season water limitations (Riddell et al. 2018). Clarifying the dominant mechanism—or combination of mechanisms—limiting distribution will improve the accuracy of forecasts and help pinpoint where habitat loss is most likely across the Appalachian region.

Disentangling plasticity from local adaptation is critical to understanding how metabolic variation arises across geographic clines (Gienapp et al., 2008; Merilä & Hendry, 2014). In our study, metabolic differences between populations were consistent with countergradient variation (Conover & Schultz, 1995): under standardized laboratory conditions, salamanders from the warmer, low-elevation environment had lower metabolic rates than salamanders from the colder, high-elevation environment. Countergradient variation occurs when local genetic adaptation offsets the direct environmental effect on phenotypic expression (Conover et al., 2009) such that apparent phenotypic similarity across environments may obscure underlying genetic divergence. Although our results align with this expectation, fully demonstrating local adaptation would require a multigenerational experiment to rule out developmental plasticity or maternal effects. Nevertheless, our *in silico* simulations show that salamanders across elevations expend similar amounts of energy in their respective environments, indicating that energy costs are not solely driven by variation in environmental temperature. Notably, we did not observe a season × elevation interaction, suggesting that metabolic compensation during winter is relatively consistent across elevations. This implies that the metabolic compensation we measured might be a relatively fixed trait, limiting the ability of salamanders to adjust to warming conditions such as those projected under climate change.

Our experimental design, which measured metabolic rates across a range of ecologically relevant temperatures, enabled us to distinguish between metabolic compensation and suppression—two fundamentally different strategies for coping with temperature. We identified compensation based on increased metabolic rates at low temperatures and suppression based on reduced metabolic rates at higher temperatures (Conover et al. 2009, White et al. 2011). Crucially, this distinction is challenging to identify from measurements at a single temperature, which may conflate the two responses. Our study design provides clear evidence of compensation in high-elevation salamanders, as they showed no suppression at warm temperatures. In the seasonal comparisons, we observed compensation at cold temperatures in winter and suppression at warm temperatures in fall. This suggests that without compensation, metabolic rates would become too low in winter to meet basic homeostatic demands, and without suppression, metabolic rate would be too high to balance energy budgets in warmer months(Fitzpatrick & Brown, 1975). Similar patterns of suppression and compensation have been documented in response to short-term acclimation experiments or seasonal temperature changes in other amphibians (Feder, 1978; Fitzpatrick, 1973; Fitzpatrick & Brown, 1975; Kiss et al., 2009), indicating that using these opposing strategies to balance homeostatic capacity and energy use may be common among amphibians. However, it remains unclear whether salamanders can dynamically employ these strategies to minimize the effect of warming climates.

Woodland salamanders have narrow environmental niches that restrict them to cool and moist habitats (Kozak & Wiens, 2006). Such species are generally considered among the most sensitive to future climate change (Botts et al., 2013; Grinder & Wiens, 2023; Thuiller et al., 2005), especially given their limited dispersal ability (Kozak & Wiens, 2010b; Liebgold et al., 2011). Woodland salamanders have long been thought to minimize energy expenditure as a core ecological strategy for coping with harsh environments (Feder, 1983; Full, 1986), with lunglessness representing a key adaptation for energy conservation (Reagan & Verrell, 1991). Nevertheless, our findings suggest that winter temperatures push salamanders to the edge of their energetic limits, forcing increases in metabolic rate to maintain basic homeostatic functions during brumation. Increasing metabolic rate at a time of inactivity and food restriction suggests that salamanders are operating near the minimum energetic thresholds compatible with vertebrate life. This study system provides a striking example of evolutionary specialization; one that may be especially vulnerable to climate change. To date, whether amphibians benefit or suffer from warmer winters remains uncertain. Some studies report improved overwinter survival and condition under milder temperatures (e.g., (Anholt et al., 2003; McCaffery & Maxell, 2010; Scherer et al., 2008; Üveges et al., 2016)) while others show the opposite—reduced survival, poor condition, or disrupted physiological processes (Reading 2010). Our results suggest that seasonal variation in the thermal sensitivity of metabolic rate, which may currently support survival under cold conditions, could become maladaptive under future warming by increasing energy demands when reserves are already limited. More broadly, our results highlight winter physiology as an underappreciated axis of climate vulnerability (Williams et al. 2015) and suggest that accurate forecasts of species persistence will require integrating seasonal metabolic dynamics alongside measurements of activity, survival, and reproduction.

## Methods

### Study system and experimental design

We collected all salamanders (*Plethodon metcalfi*, *N* = 104) in September 2023 in the temperate rain forests of the Balsam Mountain Range in the Nantahala National Forest (35°22’09.8"N 83°02’59.1"W), with authorization from the USDA Forest Service and the North Carolina Wildlife Resources Commission (ID#: 35563047). We collected salamanders from two populations, a low elevation population (∼1200 m) and a high elevation population (∼1600 m). Upon capture by hand, we placed salamanders in individual bags with locally sourced moist leaf litter and transported them to the Highlands Biological Station, where we measured body mass of each individual to the nearest 0.001 g. We then divided each population into three seasonal groups, fall (*n* = 42), winter (*n* = 41), and spring (*n* = 21). Due to sampling limitations, low elevation individuals were only represented in the fall and winter groups, and not spring (fall: *n* = 21 for low and *n* = 21 for high; winter: *n* = 21 for low and *n* = 20 for high; spring: *n* = 21 for high). Therefore, we conducted separate analyses across groups to specifically test hypotheses related to the effects of season and elevation. We then transported the fall salamanders in a cooler containing ice packs wrapped in cotton towels to the University of North Carolina at Chapel Hill the following day for metabolic measurements (see below). The winter and spring salamanders were returned to the field as part of the overwintering treatment. This study adhered to the ethical guidelines of the Institutional Animal Care and Use Committee of the University of North Carolina at Chapel Hill (IACUC ID: 23-183).

The day after capture (September 28^th^, 2023), we placed the winter and spring salamanders in underground enclosures to simulate natural winter conditions. We overwintered salamanders at the same site that we collected the high elevation salamanders (1582 m). Therefore, high elevation salamanders were kept at their native elevation, and low elevation salamanders were transplanted to a higher elevation. We designed the experiment to compare population-level differences in metabolic rate while controlling for environmental conditions. We were unable to conduct a reciprocal transplant experiment due to limits on sample size; thus we cannot make conclusions on whether differences in metabolic rate are dependent upon the elevation of the overwintering site. We also cannot determine whether differences are due to genetic or plastic effects, although a reciprocal transplant design would not provide this insight either as the experiment investigated adults (i.e., not controlling for developmental plasticity or maternal effects). Despite being limited in determining the cause of metabolic differences, our design can still make relative comparisons between high and low populations, draw inferences on whether differences are due to metabolic compensation or suppression, and use the metabolic rate data to understand the spatial and energetic consequences of winter dormancy.

Enclosures consisted of horizontal PVC pipes (14 × 5.08 cm; volume *c.* 283 mL) filled approximately halfway with locally sourced moist soil and capped on both ends with window screening mesh and zip-ties. The gauge of the window screen mesh was approximately 1.27 mm, thus preventing invertebrates from entering the enclosure while also allowing air to easily flow. To minimize environmental variation experienced by the enclosures, we maintained all enclosures within a small area (< 20 m^2^). Within this area, we manually excavated six holes approximately 60 cm × 60 cm × 30 cm deep, and we randomly assigned each enclosure to one of these six holes with respect to elevation and body mass (11 or 12 enclosures per hole in a single horizontal layer (Fig. S1). In each hole, we also placed a HOBO datalogger (HOBO U23 Pro V2 External Temperature Data Logger, HOBO, Bourne, MA 02532) that tracked date, time, relative humidity, and temperature every 30 minutes throughout the winter and spring. We note that environmental conditions did not differ between holes (see below), all salamanders survived the experiment, and we found no significant differences in mass lost between holes (for winter, *p* = 0.09; for spring, *p* = 0.68; Fig. S2). These results suggest holes are unlikely to confound our insight into the effects of physiological responses to winter. We revisited the site to retrieve salamanders for metabolic data collection on February 6^th^ and April 13^th^, 2024, for winter and spring samples, respectively. Upon arrival at the laboratory, we collected body mass from all individuals.

### Flow-through respirometry

We measured the thermal sensitivity of the volume of carbon dioxide production (V CO_2_, μL h^-1^) using a flow-through respirometry system (Sable Systems Int. [SSI], Las Vegas, NV) in a laboratory setting. While in the laboratory, salamanders were kept in individual enclosures with paper towels moistened with commercially available spring water. Due to limitations of the respirometry system, we measured salamanders over a period of three days across multiple temperatures, and salamanders rested and reached the same baseline fully rehydrated mass prior to each measurement. For the fall measurements, we ensured that physiological measurements were taken in a postabsorptive state by maintaining salamanders in an incubator (Panasonic; MIR-154) at 15°C for 15 days. For the winter measurements, salamanders had remained in the field enclosures for 131 days without food resources and were assumed to be in a postabsorptive state upon retrieval. Therefore, we initiated respirometry measurements the day after retrieving the salamanders from the enclosures to minimize the likelihood of arousal from dormancy. When not being measured, we maintained winter salamanders at 5°C between measurements to maintain salamanders at an ecologically relevant temperature that they experience underground during the winter (Fig. S2). For the spring measurements, we maintained salamanders at 15°C prior to measurements and also assumed salamanders were in a postabsorptive state after being underground for 198 days. We acknowledge that the winter salamanders experienced a different resting temperature (5°C) in the laboratory than the fall and spring populations (15°C), but this design was selected to minimize arousal from winter inactivity and improve our chances of accurately characterizing the physiological state of each seasonal group. We also note that 15°C is an ecologically relevant temperature and corresponds to nighttime air temperatures that salamanders experience in the fall and spring. Regardless, all groups experienced the same temperatures during metabolic measurements.

To assess seasonal and population-level differences in the thermal sensitivity of metabolic rates, we measured mean V CO for each individual at three ecologically relevant temperatures: 10°C, 15°C, and 20°C. Underground temperatures during winter ranged from 1– 10°C (Fig. S2), confirming that our coldest experimental condition reflected winter conditions.

In contrast, 15°C and 20°C are more representative of temperatures experienced during the active season. Our flow through system was capable of measuring seven individuals at a time (and one empty chamber used for baseline); therefore, we measured three batches (up to 21 individuals) per day at a given temperature. We also adjusted the order of temperature treatments (15°C, 10°C, and 20°C) to minimize any physiological responses to temperature ramping. Thus, each salamander experienced one temperature treatment on a given day. For each group of seven individuals, we randomly assigned an individual to each chamber and cycled through each chamber consecutively with baselines between each measurement. We measured each individual three times, with each measurement being four minutes and a total measurement period of approximately three hours. We also used a video surveillance system to monitor activity and ensure salamanders were not moving during the measurement. For fall and winter salamanders, individuals were divided equally into two groups for measurements on consecutive days (*i.e.*, the first group was measured on days 1-3 and the second group on days 4-6), which we accounted for in our statistical analysis. We note that individuals were distributed randomly between groups with respect to elevation of origin and mass.

Prior to each measurement, we recorded the mass of each individual (to the nearest 0.001 g) and then placed each salamander in an individual respirometry chamber. Salamanders then rested for 20 to 45 min prior to initiating the experiment to acclimate to the respirometry chamber and equilibrate to the experimental temperature. The respirometry system pushed air with a pump (PP2; SSI) into a bubbler bottle to saturate the air with water. This saturated airstream then continued through a dewpoint generator (DG4; SSI) to control water vapor pressure. To reduce any potential effect of variable humidity on metabolic rates, we maintained a constant vapor pressure deficit (VPD) of 0.5 kPa across temperatures, thereby ensuring each individual experienced the same evaporative demand of the air regardless of temperature. Then, the airstream was separated using a flow bar (FB8; SSI), which also controlled the flow rate to 50 mL/min. The airstream then passed through acrylic chambers (8 × 3.5 cm; volume *c*. 77 mL) containing the salamanders, suspended over hardwire mesh. The mesh promoted stereotypical posture and minimized the effect of postural differences among individuals. The respirometry chambers were housed in a temperature control cabinet with a Peltier plate (SSI) controlled by a TEMP-5 control system (TEMP; SSI) that maintained the desired temperature. After the chamber, the airstream passed through a multiplexer (MUX8; SSI), which cycled through baseline and animal chambers. From a given chamber, the airstream then passed through a water vapor analyzer (RH300; SSI) and lastly through a flow meter and a CO_2_ analyzer (CA-10; SSI). Conversion of raw data outputs from the respirometry system into physiological traits was conducted in *Expedata* (v1.9.27) using standard protocols and equations (Lighton, 2018; Riddell et al., 2018). For analysis, we used the lowest metabolic rate of the three replicates and ensured that salamanders were resting from the video recordings and exhibiting flat, stable metabolic readings consistent with rest. Any readings indicative of activity (such as irregular peaks) were not included. Prior to each experiment, we used standard procedures for zeroing and spanning the RH-300 and CA-10 using research-grade pure nitrogen, the dewpoint generator, and research grade CO_2_ mixed with nitrogen.

### Statistical analyses on metabolic rate

All statistical analyses were conducted in R (v4.2.2). We analyzed population and seasonal differences in metabolic rates using mixed-effects models with the *lme4* (Bates et al., 2015) and *lmerTest* (Kuznetsova et al., 2019) packages. Due to our experimental design, we conducted two separate analyses to understand the effects of elevation and season. In the first analysis (termed the *elevation × season* analysis), we analyzed populations from high and low elevations in the fall and winter seasons because low elevation salamanders were not represented in the spring. For the dependent variable, we log-transformed V CO_2_ to meet the assumptions of normality and homoscedasticity. For the independent variables, we included elevation origin (high, low), season of collection (fall, winter), experimental temperature (10°C, 15°C, 20°C), sex, and body mass prior to each respirometry trial. We also included the interaction between season and experimental temperature as well as the interaction between elevation origin and experimental temperature to evaluate whether the thermal sensitivity of metabolic rate differed by season or elevation. Individual was included as a random effect to account for repeated measures on an individual across temperatures. We initially included batch as a random effect (*i.e.*, first and second batch of respirometry); however, variance estimates for this term were near zero (resulting in a singularity warning). Therefore, batch did not explain sufficient variation in the response variable to warrant being included in the model and was excluded to improve interpretability and avoid overparameterization.

In the second analysis (termed the *season* analysis), we analyzed seasonal variation in metabolic rate across salamanders collected in the fall, winter, and spring for individuals collected from high elevations. The model was identical to the first model, except for excluding elevation as a factor in the model and including spring as an additional factor for the effect of season. This model was designed to test metabolic differences across fall, winter, and spring salamanders and assess whether spring individuals showed metabolic rates more similar to fall individuals, indicating a physiological transition out of winter dormancy.

For both analyses, we conducted pairwise *post hoc* t-tests to evaluate differences in metabolic rate between seasonal and elevational groups at each temperature (e.g., fall vs. winter metabolic rate at 10°C). These comparisons were specified *a priori* to test biologically motivated hypotheses about metabolic compensation and suppression, and to avoid unnecessary multiple comparison corrections (*e.g.*, metabolic rate in fall at 10°C *versus* in winter at 20°C). Because our comparisons were specified *a priori* to test biologically motivated hypotheses, we did not apply corrections for multiple comparisons. However, applying a Bonferroni correction for each family of tests (adjusted significance threshold = 0.017 for season or elevation differences across experimental temperatures) does not alter the interpretation of our results.

### Statistical analyses on body mass

We also evaluated whether changes in body mass across seasons and elevations could reveal consistent patterns in energetic use during winter dormancy. In addition to evaluating differences between seasons and elevations, we also tested whether mass loss was predictable from metabolic rate, assuming the primary substrates were lipids or glycogen (see Supplementary Materials). However, despite salamanders being held without access to food in fully saturated conditions (as confirmed by HOBO dataloggers), we found that most individuals lost more mass than could plausibly be attributed to fuel metabolism alone (Fig. S3). These results suggest that mass loss was primarily driven by water loss—whether passive or physiologically mediated— rather than strictly by the catabolism of energetic reserves. As a result, we were unable to disentangle water loss from fuel metabolism and therefore could not draw reliable conclusions about energetic strategies from changes in body mass.

### Biophysical species distribution model

We used a mechanistic species distribution model (Riddell et al., 2018) to estimate winter energetic costs for *P. metcalfi* across their geographic range under current and future climate scenarios (USGS, 2018). These models blend climate data with the ecology, natural history, morphology, behavior, and physiology of salamanders to quantify axes of performance, such as energy expenditure (Briscoe et al., 2023, Kearney & Porter, 2009; Riddell et al., 2023). By downscaling macroclimatic conditions to the microhabitat conditions that salamanders experience underground during winter, we assessed how variation in metabolic rate during dormancy influences energy reserves. We aimed to evaluate whether winter energetics influence current geographic range limits and to assess how metabolic compensation during brumation could alter overwinter energetic costs under future warming scenarios.

### Environmental variables

We downloaded monthly minimum and maximum air temperature from *Worldclim* at a 30s resolution (1 km^2^) for both current (1970) and future (2081-2100; Shared Socio-economic Pathway [SSP] 245) climate conditions. For future warming scenarios, we averaged across all available global circulation models (GCMs) to produce an average warming scenario that accounts for the uncertainty across GCMs (Tebaldi and Knutti, 2007; Beaumont et al., 2008, Parker, 2013). We used SSP 245 because it represents one of the least extreme warming scenarios, with temperatures increasing by 2.1 – 3.5 by 2100 (IPCC, 2023). We estimated hourly air temperatures from minimum and maximum air temperature using standard analytical approaches (Campbell and Norman, 1998; Riddell et al., 2019a). We also estimated soil temperature by using surface balance energy equations (Leaf and Erell, 2018; Porter et al., 2023) that account for sensible heat flux, net radiative flux, ground storage of heat, and latent heat flux. To validate the soil model, we compared predicted soil temperatures to observed soil temperatures for southern Appalachia recorded by the Soil Climate Analysis Network (SCAN) across ten sites and four soil depths (2in, 4in, 8in, 20in). Because observed soil temperatures never dropped below freezing (Figures S5-8) due to the zero-curtain effect (Outcalt et al., 1990), we set predicted minimum temperatures to ≥ 1 to maximize model performance. The soil performed well at predicting soil temperautres in the Appalachians (*R^2^* = 0.92; mean slope ± error: 0.86 ± 0.001), performing as well as or better than other similar approaches (Kearney et al., 2014). Our model also estimated actual vapor pressure of the air by assuming that the minimum air temperature was the dewpoint temperature (Kimball et al., 1997; Newman et al., 2022), and saturated air pressure was estimated using equation S7 in Riddell et al. (2018). These data were then combined with the ecology and physiology of woodland salamander to predict energetic costs.

Our simulations were designed to understand the energetic consequences of metabolic responses to the winter months when salamanders are underground and inactive. Therefore, we used our soil model to calculate soil temperature 20 cm below the ground surface as an estimate of body temperature, a depth similar to our experimental design and observed in lungless salamanders (Taub 1961, Spight 1967). We ran multiple iterations of our biophysical model to estimate energetic costs that incorporated seasonal variation in metabolic rates. Specifically, we used the metabolic rates from individuals collected in the fall and winter to calculate standard metabolic rate using the following equations:

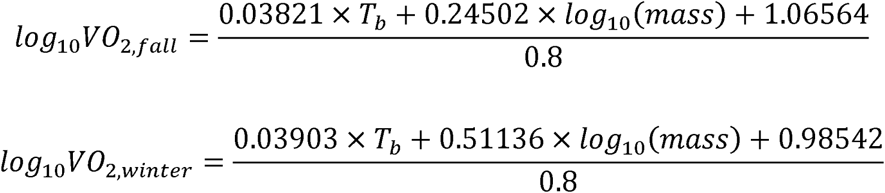

where *log*_10_*VO_2,fall_* is the log-transformed volume of oxygen consumption for salamanders during the fall, *log*_10_*VO_2,winter_* is the volume of oxygen consumption for salamanders during the winter (µL/h), *T_b_* is the body temperature of the salamander, and *mass* is the mass (g) of the salamander. These equations were derived from the final statistical models used to estimate temperature- and mass-dependent variation in metabolic rates for each season. Assuming salamanders were consuming lipids, we used a respiratory quotient of 0.7 to calculate V O_2_ from our V CO_2_ measurements (Smith 1976; Riddell et al., 2018). We then transformed V O_2_ to kJ/h by converting our estimates to mL/h, followed by multiplying the values by 20.1 J/mL of O_2_, and finally converted the total kJ of energy consumed during that period into grams of lipids by dividing energetic cost (kJ) by 37 (Zani et al., 2012). *Plethodon* are reported to have lipid reserves that make up 8-25% of their dry mass (Fitzpatrick, 1973; Fitzpatrick, 1976; Camp & Jenson, 2007). To avoid potentially over- or underestimating lipid reserves, we averaged the reported values for a final lipid reserve of 14.75% of the dry mass. Assuming salamanders consist of approximately 80% water (Crump, 1979), we estimated that an average size adult (3 g or 0.6 g dry mass) had approximately 0.0885 g of lipid reserves. We then divided the total grams of lipids lost by the grams of lipid reserves to calculate the proportion of the lipid reserves consumed for each climate scenario and thermal sensitivity of metabolic rate (fall or winter). We ran each model (*VO_2,fall_* and *VO_2,winter_*) over the inactive season (November to March) to determine how seasonal variation in metabolic rate affects energy reserves across the geographic range of the focal species. We considered sites locally extinct when salamanders’ lipid consumption surpassed available reserves (Riddell et al. 2018).

To evaluate whether energy reserves predicted the geographic distribution of our focal species, we used presence locations for *P. metcalfi* (downloaded from GBIF) to determine if presence locations were associated with proportion of lipids consumed across the landscape under the current climate scenario assuming the winter metabolic rates, the most ecologically relevant rate. We first extracted the values of the predicted proportion of lipids lost for each presence point across the landscape (*N* = 8,130). We then created a three-kilometer buffer around each presence location and removed the buffered extents from the species’ range to create a landscape of locations where the focal species have not been found. Using the remaining portion of the range, we randomly generated the same number of pseudo-absence points. We then extracted the predicted proportion of lipids consumed that corresponded to each absence. We used a logistic regression to evaluate the relationship between the proportion of lipids consumed and the presence (1) or absence (0) of our focal species. This approach allowed us to evaluate whether predicted depletion of lipid reserves during winter was associated with a near-zero probability of species presence, indicating potential energetic constraints on distribution.

### *In silico* reciprocal transplant

We tested for countergradient variation by examining whether salamanders from low elevations (with lower thermal sensitivity) had similar energy expenditure to salamanders from high elevations (with higher thermal sensitivity) at their native elevation, indicating potential adaptive compensation across environmental gradients. For the elevational models, we calculated monthly energetic cost from lab-measured metabolic rates of individuals collected at low and high elevation using the following equations:

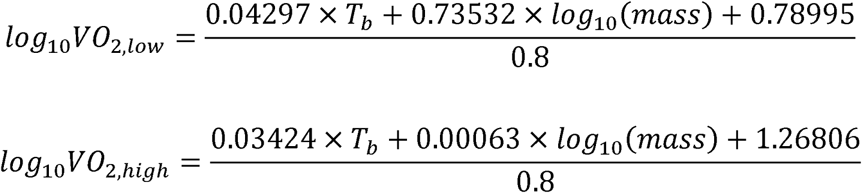

where *VO_2,low_* is the volume of oxygen consumption for low elevation species, and *VO_2,high_* is the volume of oxygen consumption for high elevation species (µL/h). We then obtained environmental temperatures for the high elevation site from the underground data loggers deployed during the winter. To approximate low elevation temperatures, we adjusted the high elevation temperature data by adding 2.5°C, based on multi-year environmental monitoring that shows low elevation near-surface temperatures average 2.5°C warmer than those at high elevation (Riddell and Sears 2015). We analyzed the differences in estimated metabolic rates using a two-way ANOVA with elevation and thermal sensitivity as factors. We performed post hoc pairwise comparisons and calculated effect sizes using the eff_size() function from the emmeans package, which provides standardized mean differences (Cohen’s d) to quantify the magnitude of differences between groups.

### Histological analyses

To assess histological indicators of brumation, we dissected tissue from the dorsal skin and lower intestine. We quantified seasonal variation in epidermal thickness, intestinal diameter, and number of villus foldings, as variation in traits associated with these tissues has been identified as markers of dormancy in amphibians (Barni et al., 1999, 2002; do Nascimento et al., 2016; Hume et al., 2002). We euthanized salamanders by immersion in MS-222 solution (0.5 – 2.0 mg per L) buffered with sodium hydroxide to balance pH until they were unresponsive (absence of righting reflex), followed by secondary euthanasia via decapitation. We removed the dorsal skin and lower intestinal tissue, which were immediately immersed in a 10% formalin solution. After seven days, we transferred tissues to a 70% ethanol solution for storage prior to processing. Histological sectioning and staining were performed by the Histology Research Core Facility at the University of North Carolina at Chapel Hill. Tissues were embedded in paraffin, sectioned transversely at 5 µm, and stained using a standard hematoxylin and eosin (H&E) protocol.

We used ImageJ (v1.54) to quantify morphological variation in skin and intestinal tissue across seasons. For dorsal skin, we obtained 2 to 12 sections from each of 22 individuals (8 fall, 8 winter, 6 spring). For each section, we captured two non-overlapping images at 10× magnification and used the linear measurement tool to measure epidermal thickness at 30 equidistant points per image. For the lower intestine, we obtained 2 to 10 sections per individual from 21 salamanders (7 fall, 8 winter, 6 spring). Using the same the linear measurement tool, we recorded 15 diameter measurements per section, taken from 15 evenly spaced angles across the cross-section of the intestine. Additionally, we measured villus length by drawing a straight line from the base of the lumen to the peak of each villus (Secor, 2005), and in doing so, we also recorded the total number of intestinal villi in each section.

We analyzed the epidermal thickness, intestinal diameter, villus length, and villus number using linear models or linear mixed effects models. For villus length, intestinal diameter, and epidermis thickness, we used mixed-effects models to accommodate repeated measures of individuals and included individual identity as a random effect, body mass as a covariate, and season as a factor. In contrast, villus number showed no within-individual variation except for one salamander. Therefore, for villus number, we used a simple linear model with season as a fixed effect and body mass as a covariate, using the median villus count for the one individual with variable measurements. We then used a similar *post hoc* statistical framework as the metabolic rate analyses for comparing seasonal differences in histological measures.

## Data availability

All data are available through the Open Science Framework at https://osf.io/2bwj9/. All scripts are available at https://github.com/braulioassis/Plethodon-dormancy.

## Acknowledgements

We thank the Highlands Biological Station for logistical support during collection.

