## Supplementary information for "Winter metabolic compensation endangers woodland salamanders under climate change"

### Supplemental Methods

### *Analysis for mass loss during dormancy*

We investigated whether individual variation in mass loss between fall and winter were associated with metabolic rates or elevation of origin. As in our primary analyses on metabolic rate, we used two separate models: one comparing mass loss between low- and high-elevation salamanders from fall to winter, and another comparing mass loss across seasons (fall to winter and fall to spring) in high-elevation salamanders only. In the first model, we used proportion mass loss (expressed as the change in mass divided by the initial mass) as the response variable and elevation of origin (low or high), sex, and log-scaled *V̇*CO_2_ measured at 10ºC as predictors. We used metabolic rate at 10ºC because this temperature was the most representative of the temperatures experienced during winter and might explain variation in mass loss if driven by energy consumption. We also included a random effect for the hole in which a salamander overwintered. We did not include body mass as a predictor to avoid issues related to regression to the mean (Gunderson, 2023), and we note that using the change in mass with mass as a covariate provided qualitatively similar results. In the second analysis across all three seasons, we determined whether salamanders showed differences in mass lost between fall and winter and between fall and spring, using the same modeling framework but with season (winter or spring) as a fixed effect. For both analyses, we conducted Type II Analysis of Covariance (ANCOVA) to assess significance.

### *Estimating lipid mass loss during winter dormancy*

We used temperature data from underground HOBO dataloggers to estimate energy expenditure between seasonal time points and elevations, applying the metabolic rate equations described in the main Methods section. Specifically, to estimate energy expenditure, we first converted volume of carbon dioxide to grams of lipids consumed per hour using the same assumptions described in the main text. We then used a linear mixed-effects model that related log-transformed lipid depletion rates to body mass, experimental temperature, and season, with individual identity included as a random effect. We generated two temperature time series from HOBO dataloggers deployed underground at field sites and paired these with representative seasonal body masses (~2.6 g) to estimate lipid consumption over winter and spring time spans. We used these model predictions to estimate hourly lipid use across the time series, then integrated these values to calculate total lipid depletion for each season and elevation. Predictions were back-transformed from the log scale, and we calculated prediction intervals using ±3 standard deviations of the fitted values, representing a 99.7% interval under the assumption of normally distributed errors. This wide interval captures nearly all expected variation in predicted lipid use, providing the upper and lower bounds of physiological energy expenditure. For lipid loss across seasons, we estimated the mass lost between fall and winter as well as fall and spring for high elevation salamanders, due to the lack of low elevation salamanders in the spring sample. For lipid mass loss across elevations, we estimated the mass lost between fall and winter for high and low elevation salamanders. We acknowledge that some ectotherms catabolize glycogen during brumation (Peng et al. 2021); however, prior studies in woodland salamanders (e.g., Plethodon spp.) demonstrate consistent lipid depletion across winter dormancy (e.g., Camp & Jensen, 2007; Fraser, 1976), providing strong evidence that lipid catabolism is the primary energetic pathway in this group. As such, we focused our analyses on lipid loss as the most biologically relevant measure of overwintering energy expenditure.

**Supplemental Results**

*Mass loss during dormancy*

Salamanders from high elevations lost more mass than salamanders from low elevations between fall and winter (F_1,37_ = 6.38, *P* = 0.016; Fig. S4). Specifically, low elevation salamanders lost 5.0% of their body mass whereas high elevation salamanders lost 10.4% (Fig. 2). Mass lost was not associated with *V̇*CO_2_ (*P* = 0.55) nor sex (*P* = 0.46). For the analysis across seasons, salamanders lost similar proportions of mass from fall to winter and from fall to spring (*P* = 0.20). However, we note that salamanders lost slightly more mass on average from fall to spring (12.0%) than fall to winter (10.5%), suggesting most of the mass loss occurred from the fall to the winter. We did not observe any significant effects of sex (*P* = 0.72) or *V̇*CO_2_ (*P* = 0.38) on mass loss between seasons.

*Lipid mass loss during winter dormancy*

Our estimates of mass loss indicated that salamanders lost substantially more mass than would be expected from the metabolism of lipids alone. From fall to winter, low-elevation salamanders lost approximately 5.0% of their body mass, compared to predicted losses of 2.4% if relying solely on lipid stores (Fig. S4). High-elevation salamanders lost approximately 9.0% of their body mass, while predicted losses were only 3.1%. For high elevation salamanders, individuals lost 7.0% of their body mass from fall to winter and 12.0% from fall to spring. In contrast, salamanders were predicted to lose only ~3.1% for those same periods, assuming metabolism of lipids. Given this large discrepancy between observed and predicted values, it is likely that most of the mass loss was due to water loss rather than fuel consumption, although we were unable to confirm this directly.

### Supplemental Figures

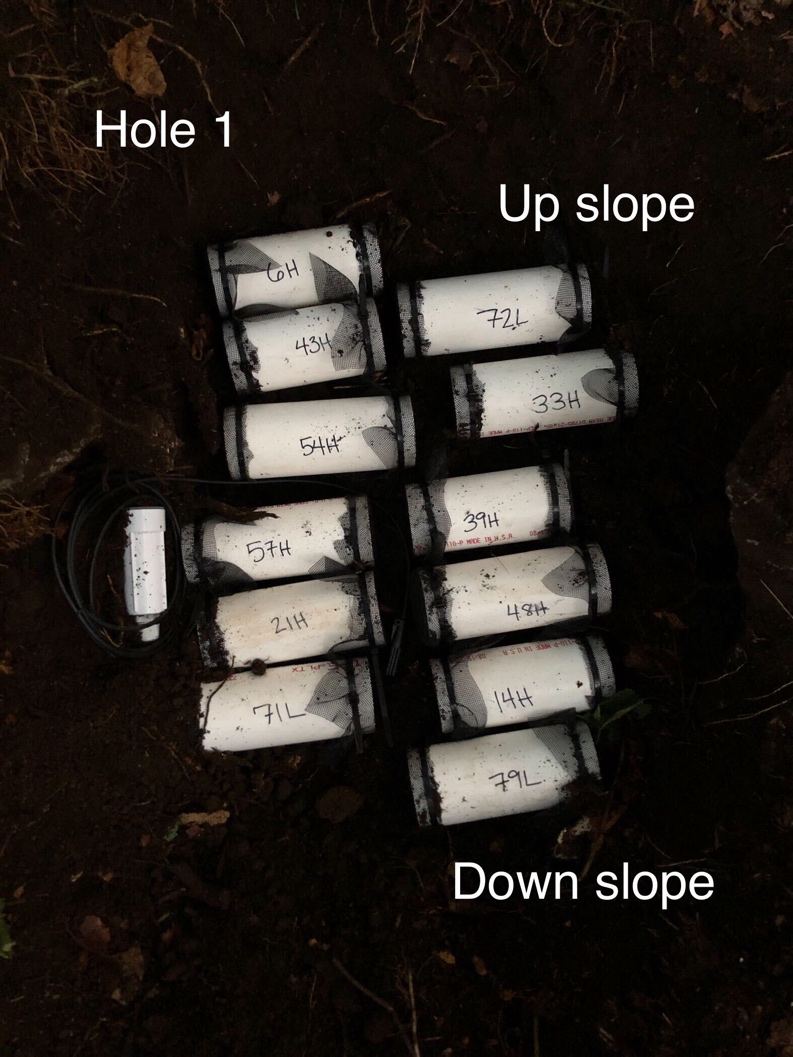

**Figure S1:** **An example of the individual enclosures and holes for the overwintering field experiment**. Enclosures placed in one of six total holes housing individual salamanders from September 28^th^, 2023 until either February 6^th^ (winter salamanders) or April 13^th^, 2024 (spring salamanders). Enclosures were buried in holes approximately 60 × 60 × 30 cm deep across a 20 m^2^ area at 1582 m above sea level.

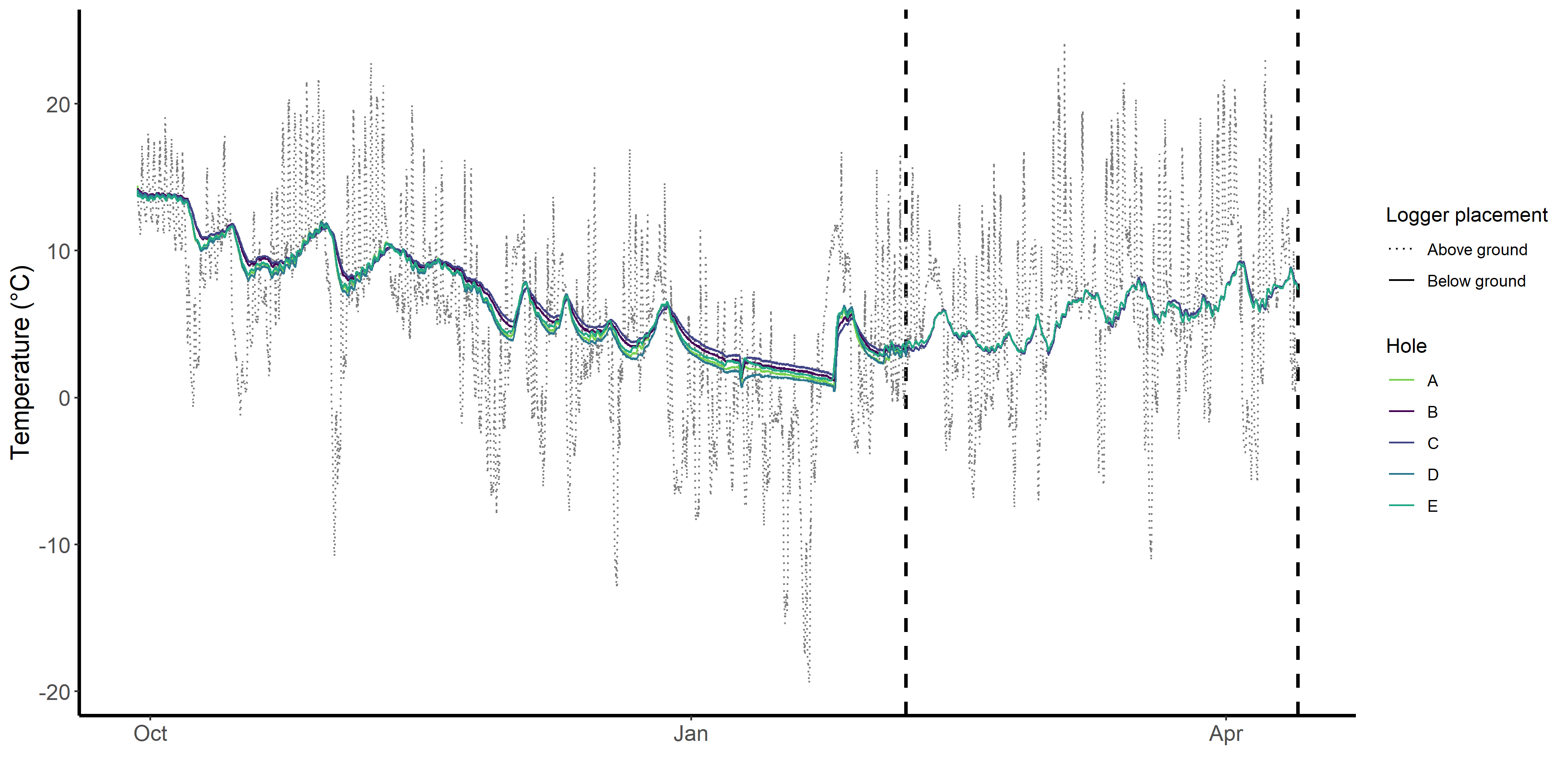

**Figure S2**: **Environmental temperatures throughout the dormancy experiment**. Dotted line corresponds to the average temperature recorded by two loggers hung approximately 160 cm above ground on trees in the vicinity of the dormancy site. Solid colored lines correspond to temperatures registered by loggers placed alongside the salamander enclosures approximately 30 cm below ground. Dashed vertical lines correspond to the winter and spring time points upon which salamanders were retrieved for metabolic and histological analyses. Note that half of the loggers were removed at the first vertical dashed line when winter salamanders were collected.

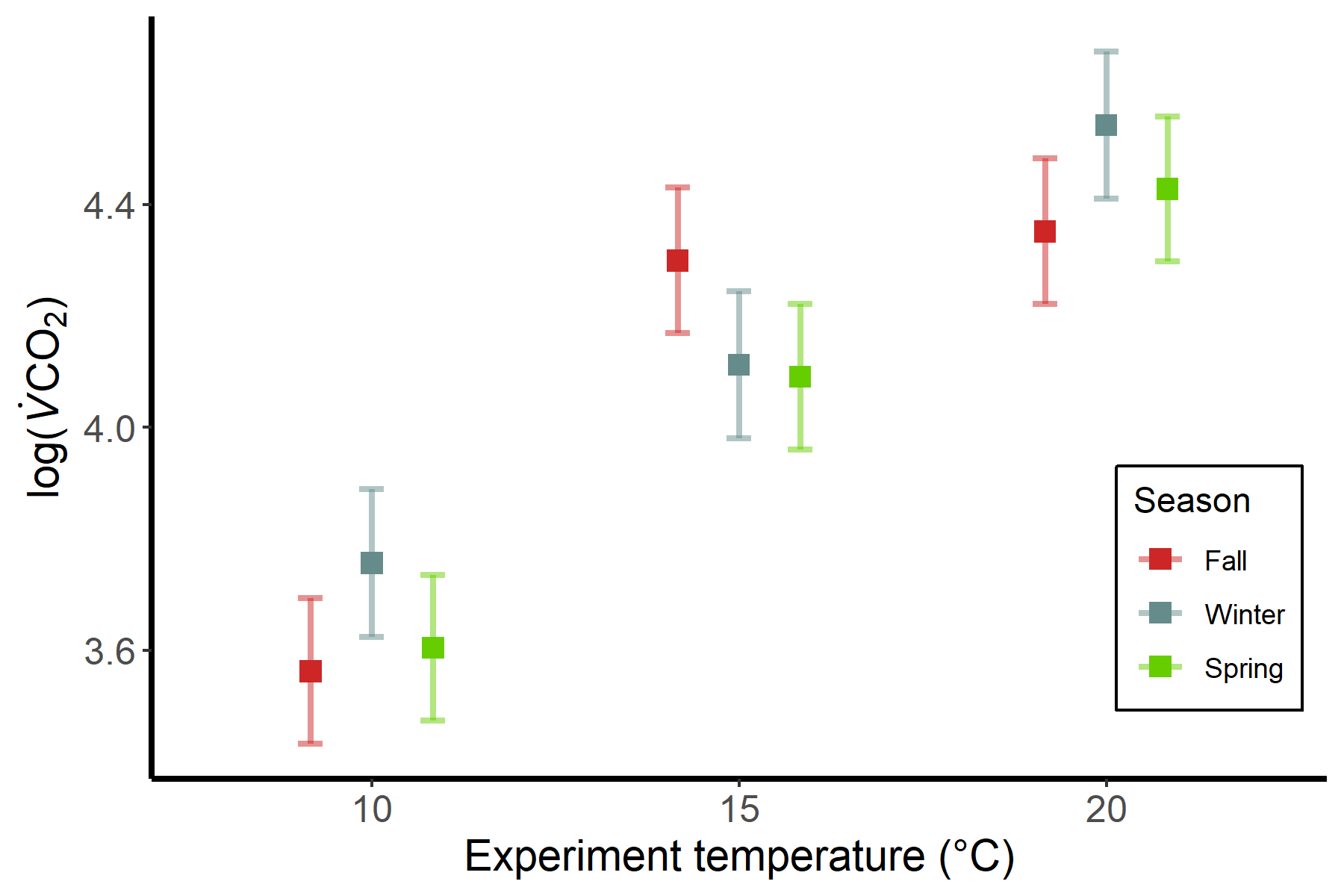

**Figure S3:** **Experimental temperature and season influenced variation in metabolic rate upon including salamanders collected in the spring**. The figure shows a significant interaction between season and experimental temperature on log-transformed (*F* = 3.47, *P* = 0.010). Points represent marginal means with 95% confidence intervals for a season-by-temperature interaction term controlling for the effects of sex and body mass, along with individual identity as a random effect. Low elevation salamanders were excluded from the analysis because they were not represented in the spring sample.

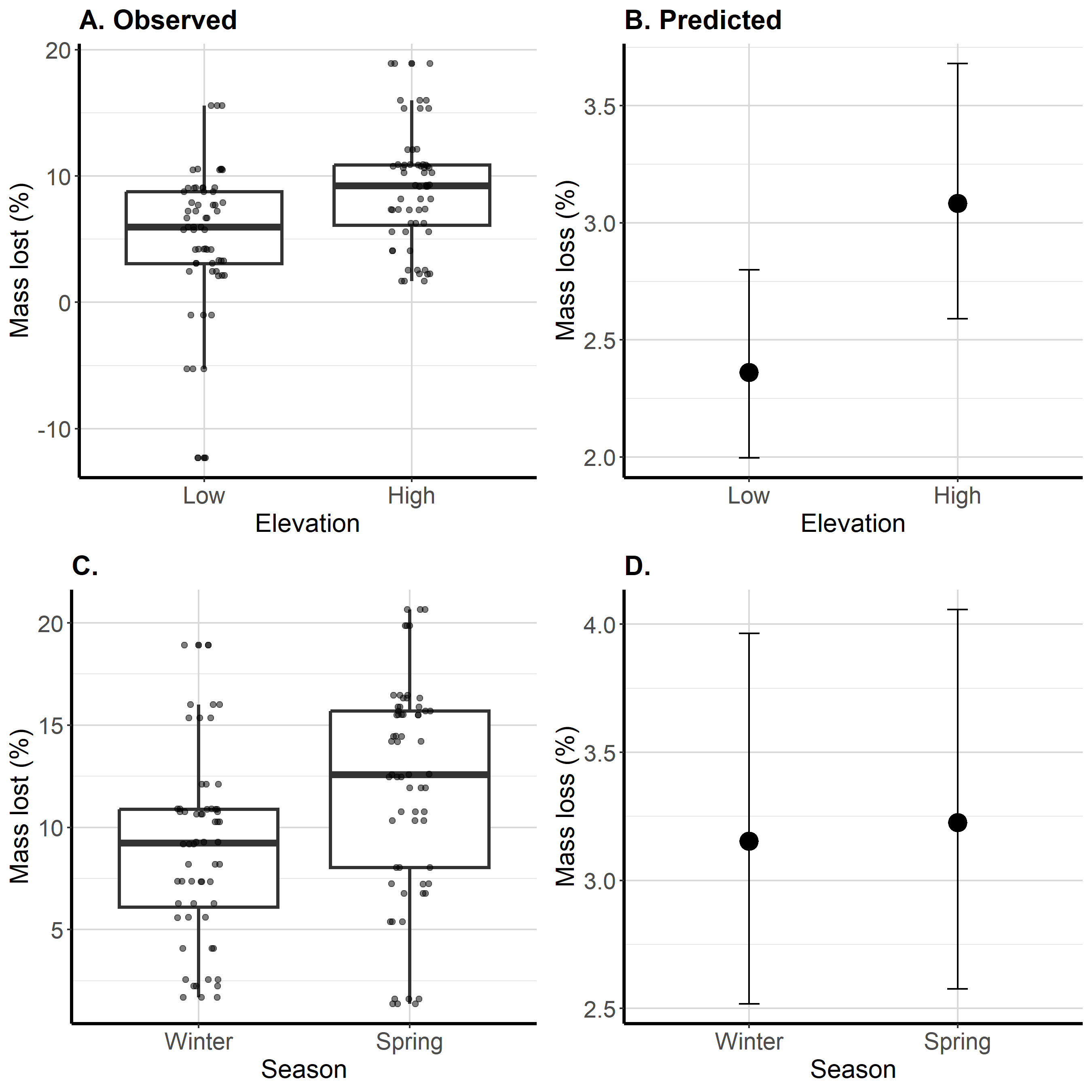

**Figure S4:** **Observed and predicted mass loss throughout the experiment**. (A) Salamanders from low elevation lost significantly less mass between fall and winter than high elevation salamanders (F_1,37_ = 6.38, *P* = 0.016). Boxplots illustrate the 25^th^, 50^th^, and 75^th^ percentiles, and whiskers represent the highest and lowest values within 1.5 times the interquartile range. (B) Predicted percent mass loss based on population-specific metabolic rates and experienced soil temperatures. Mean percent lipid loss ± 99.7% prediction intervals are shown. (C) Salamanders lost significantly more mass between fall and spring than between fall and winter (F_1,37_ = 4.16, *P* = 0.049). (D) Predicted percent mass lost between fall and winter as well as fall and spring based on experienced ambient temperatures. Mean percent lipid loss ± 99.7% prediction intervals are shown.

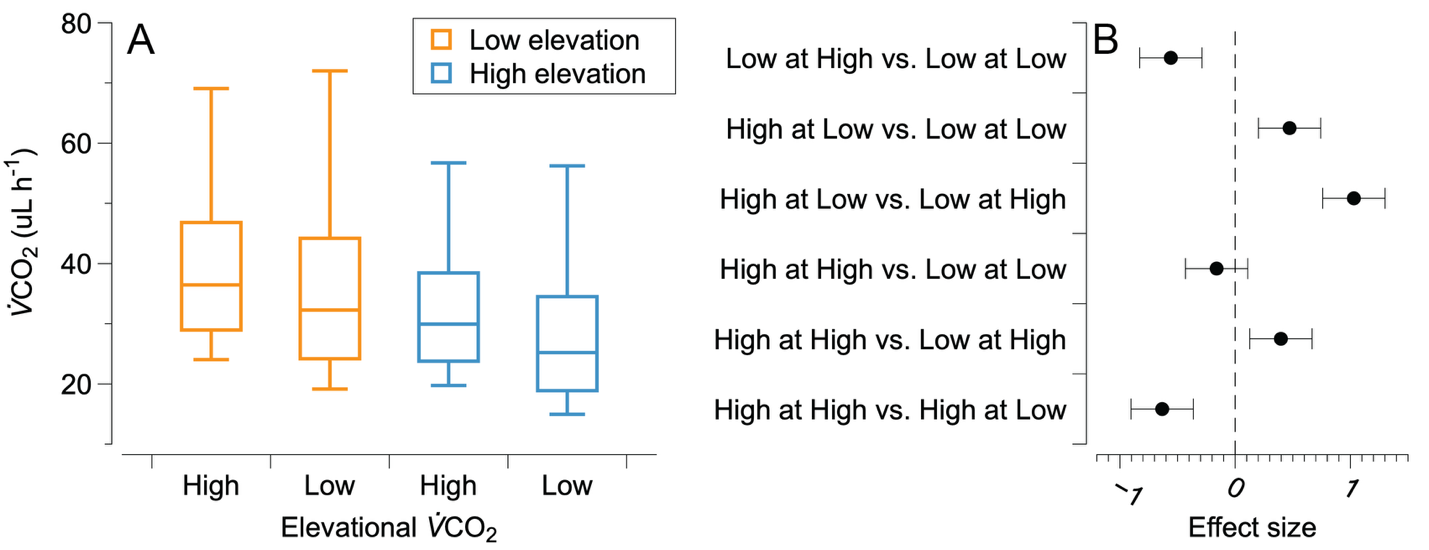

**Figure S5:** **Testing for countergradient variation in metabolic rate**. (A) Average *V̇*CO_2_ for simulated reciprocal transplants based on population-specific thermal sensitivities of metabolic rate and thermal environments. Box plots are shown for the low elevation site (orange) and high elevation site (blue) with a thermal sensitivity for metabolic rate at high or low elevations. (B) Effect sizes for pairwise comparisons between all categories of simulated energetic costs. Comparisons are made between, for instance, salamanders with low elevation metabolic rates simulated at high elevations are compared with salamanders with low elevation metabolic rates at low elevations (“Low at High vs. Low at Low”). The effect sizes on energetic costs between population-specific thermal sensitivities within their native thermal environments overlap zero (that is, “High at High vs. Low at Low”), suggesting that metabolic compensation results in countergradient variation for metabolic rates across elevations.

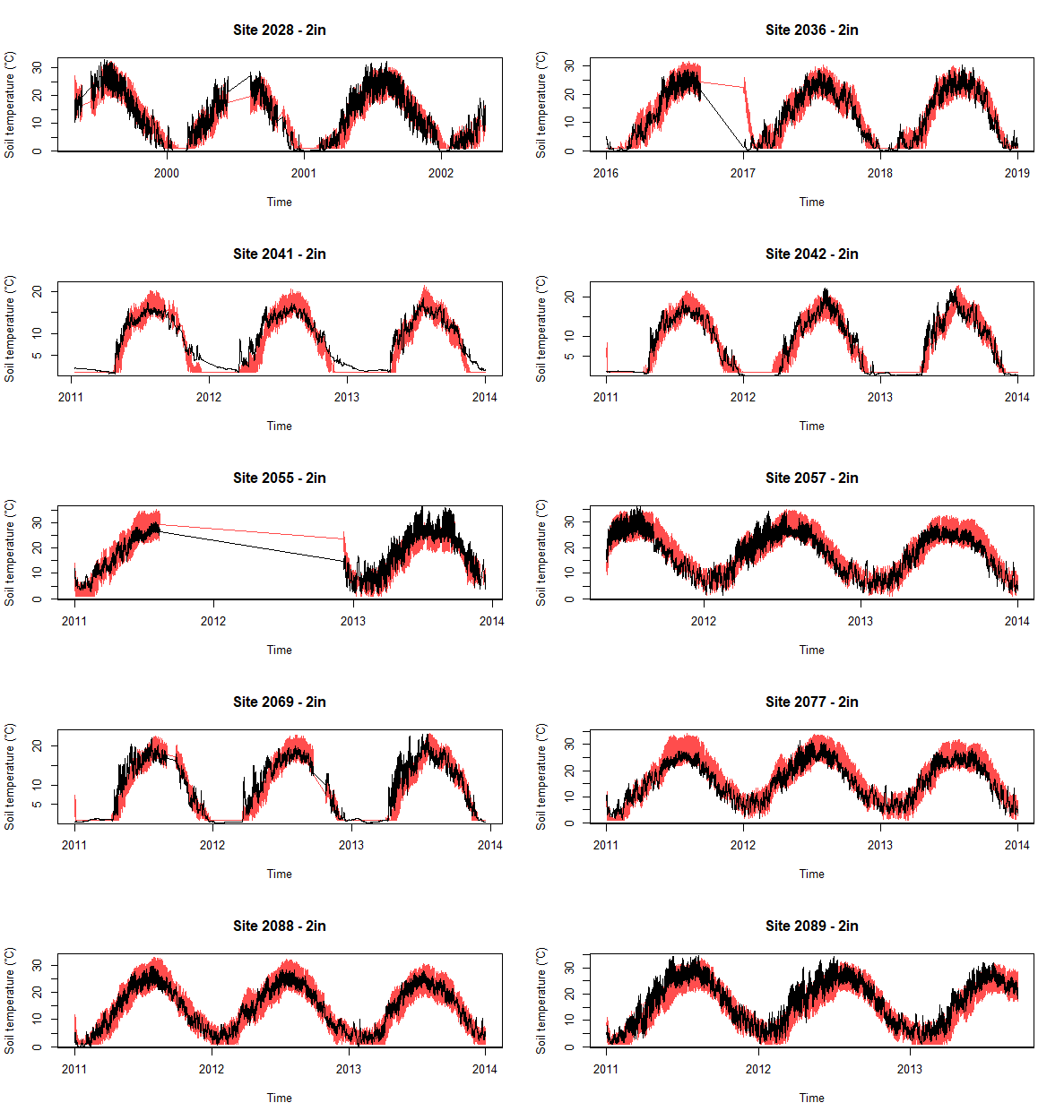

**Figure S6: Validation of soil temperature model at 2 in.** Output of soil validation model for a depth of 2 in across 10 sites. Black line is observed soil temperatures, and red line is predicted soil temperatures. Assuming minimum temperatures of ≥ 1℃, average *R^2^* across sites and soil depths is 0.92 and mean slope ± standard error = 0.86 ± 0.002. Straight lines connecting data are indicative of missing data.

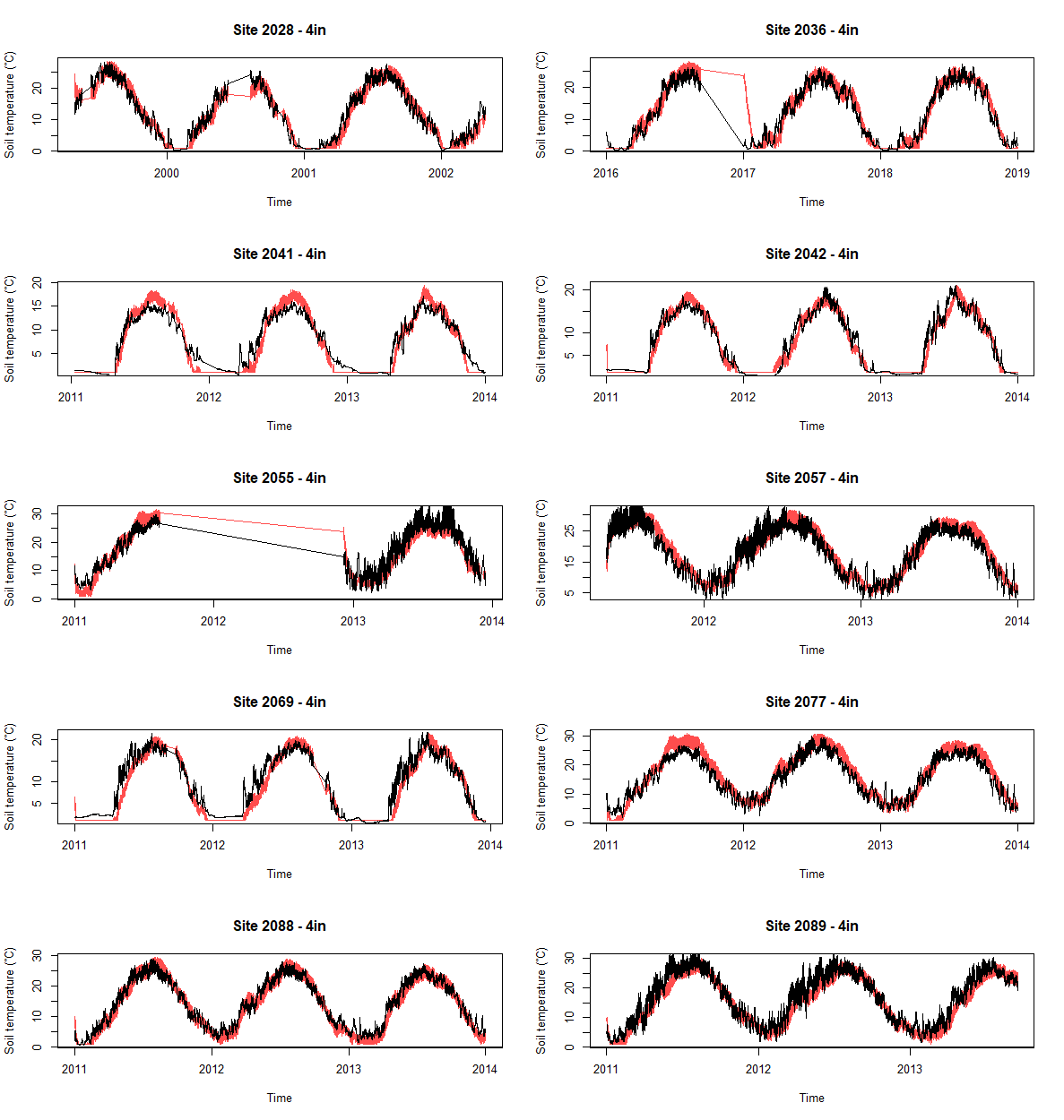

**Figure S7: Validation of soil temperature model at 4 in.** Output of soil validation model for a depth of 4 in across 10 sites. Black line is observed soil temperatures, and red line is predicted soil temperatures. Assuming minimum temperatures of ≥ 1℃, average *R^2^* across sites and soil depths is 0.92 and mean slope ± standard error = 0.86 ± 0.002. Straight lines connecting data are indicative of missing data.

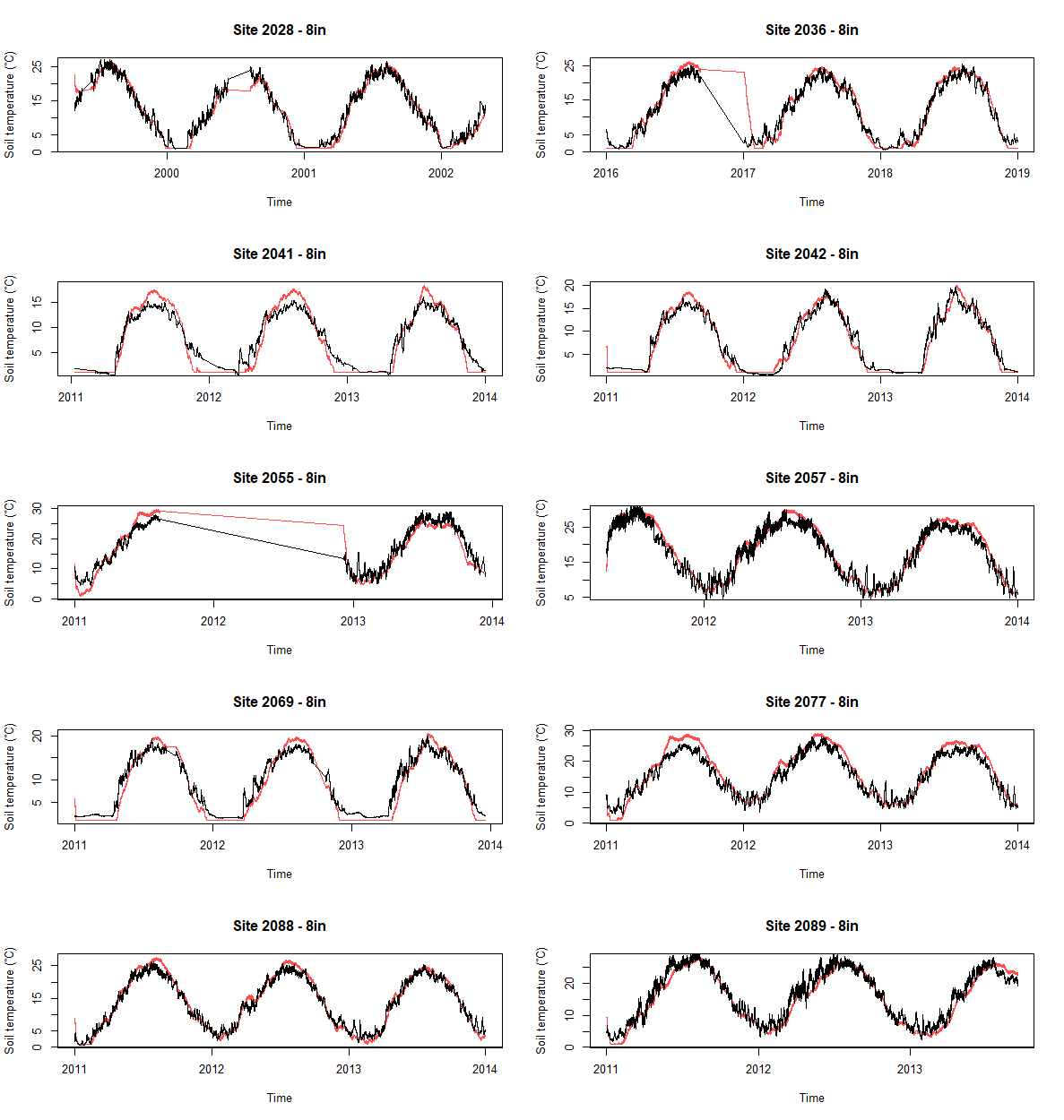

**Figure S8: Validation of soil temperature model at 8 in.** Output of soil validation model for a depth of 8 in across 10 sites. Black line is observed soil temperatures, and red line is predicted soil temperatures. Assuming minimum temperatures of ≥ 1℃, average *R^2^* across sites and soil depths is 0.92 and mean slope ± standard error = 0.86 ± 0.002. Straight lines connecting data are indicative of missing data.

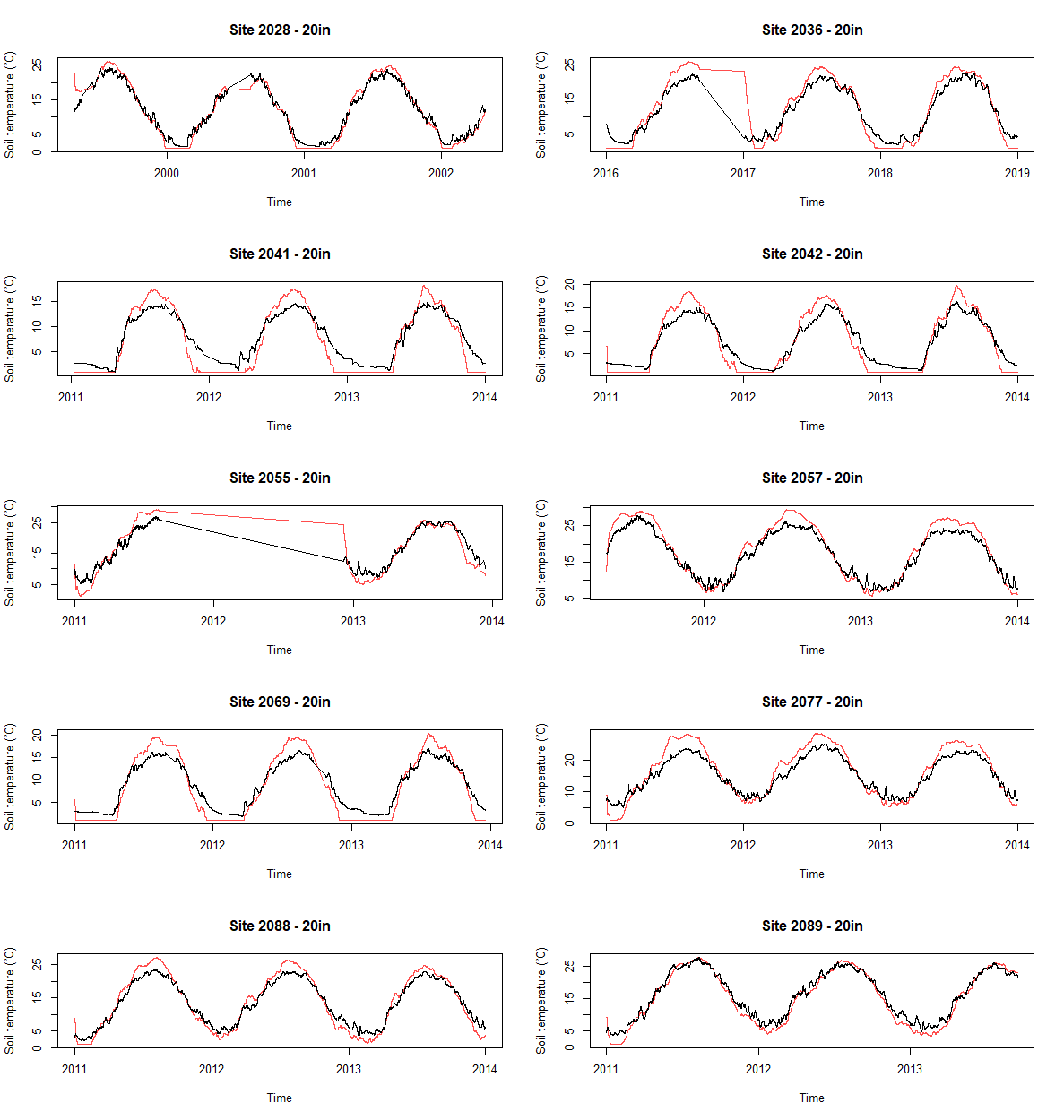

**Figure S9: Validation of soil temperature model at 20 in.** Output of soil validation model for a depth of 20 in across 10 sites. Black line is observed soil temperatures, and red line is predicted soil temperatures. Assuming minimum temperatures of ≥ 1℃, average *R^2^* across sites and soil depths is 0.92 and mean slope ± standard error = 0.86 ± 0.002. Straight lines connecting data are indicative of missing data.
